# Hominini-Specific Regulation of *CBLN2* Increases Prefrontal Synaptogenesis

**DOI:** 10.1101/2019.12.31.891069

**Authors:** Mikihito Shibata, Kartik Pattabiraman, Sydney K. Muchnik, Nenad Sestan

## Abstract

The similarities and differences between nervous systems of various species result from developmental constraints and specific adaptations ^1-4^. Comparative analyses of the prefrontal cortex (PFC), a region of the cerebral cortex involved in higher-order cognition and complex social behaviors, have identified confirmed and putative human-specific structural and molecular changes ^4-8^. For example, one crucial specialization involves the anterior-posterior gradient in synaptic density, with a disproportionately higher number of dendritic spines specifically in the human PFC compared to other analyzed primates ^5^. These changes are likely mediated by divergence in spatio-temporal patterns of gene expression ^9-17^, which are prominent in the mid-fetal human cerebral neocortex ^15,18-20^. Analyzing developmental human and macaque brain transcriptomic data ^15,20^, we identified a transient PFC- and laminar-specific upregulation of the gene encoding cerebellin 2 (CBLN2), a neurexin (NRXN) and glutamate receptor delta (GRID/GluD)-associated synaptic organizer ^21-27^, in human mid-fetal development coinciding with the initiation of synaptogenesis. Moreover, we show that this difference in expression level and laminar distribution of *CBLN2*, is due to Hominini-specific deletions affecting SOX5 binding sites within a retinoic acid-responsive *CBLN2* enhancer. *In situ* genetic humanization of the mouse *Cbln2* enhancer drives increased and ectopic laminar *Cbln2* expression and promotes glutamatergic and GABAergic synaptogenesis specifically in the PFC. These findings identify a putative genetic and molecular basis for the disproportionately increased connectivity in the Hominini PFC and suggest a developmental mechanism linking dysfunction of the NRXN-GRID-CBLN2 complex, to the pathogenesis of neuropsychiatric disorders.

## Introduction

Expansion of the PFC in Old World anthropoid primates (Catarrhini) is associated with derived characteristics such as increased dendritic complexity and synaptic spines of the glutamatergic excitatory projection (pyramidal) neurons compared to more posterior cortical association and sensory areas, creating an anterior-posterior gradient of synaptic density ^28-30^. Moreover, this gradient is exaggerated in humans by the disproportionately high number of spines on pyramidal neurons in the human PFC, as compared to other analyzed primates ^5^. The anterior-posterior gradient of early neocortical synaptogenesis begins during the human mid-fetal period or equivalent developmental age in other mammals^7,31^, before proceeding in an anterior-posterior direction ^32^ and continues at an accelerated rate well beyond birth ^33,34^. Identifying the genetic basis and molecular mechanisms underlying this anterior-posterior gradient of synaptogenesis may provide clues to the origin of increased synaptic density within the human PFC and some aspects of human uniqueness, cognition, and neuropsychiatric disease. In our accompanying study ^35^, we identified a prefrontal-enriched gradient of retinoic acid (RA) concentration and pattern of gene expression in the human mid-fetal neocortex. Among the identified genes were those linked to synaptogenesis, including *CBLN2*, which had the greatest loading of differentially expressed genes representing the anterior-posterior neocortical axis, and was upregulated in the mid-fetal PFC compared to the prospective primary motor cortex (M1C) and other non-PFC areas ^35^. Thus, we hypothesize that RA-mediated regulation of *CBLN2* expression in the human mid-fetal PFC underlies the disproportionate number of synapses in the PFC.

### Species-specific enrichment and laminar pattern of *CBLN2* expression in mid-fetal PFC

Members of the CBLN family encode secreted neuronal glycoproteins, which serve as excitatory and inhibitory synaptic organizing molecules ^21-27^. CBLN2, along with its binding partners, the neurexin (NRXN) family of cell adhesion proteins and the glutamate receptor delta (GRID/GluD) receptors ^23-27^, have been implicated in various neuropsychiatric diseases including autism spectrum disorders (ASD), schizophrenia, Tourette’s syndrome, and obsessive-compulsive disorder (OCD) ^36-39^. By analyzing RNA-sequencing (RNA-seq) transcriptomic data from eleven neocortical areas of the human and macaque brain across prenatal and postnatal development from the BrainSpan and PsychENCODE projects ^15,20^ (brainspan.org and evolution.psychencode.org), we identified a precocious and transient increase in human (1.8-fold change) and macaque (2.0-fold change) *CBLN2* expression in the major areas of the prospective PFC (i.e., medial, mPFC/MFC; dorsolateral, dlPFC/DFC; ventrolateral, vlPFC/VFC; and orbital, oPFC/OFC) compared to the seven analyzed non-PFC areas during mid-fetal development (13-24 postconception weeks, PCW) or periods 4-6 according to Kang et al. ^40^ (Student’s t-test, human P = 2.2e-16; macaque P = 1.7e-9; **Fig. 1a**) ^20^. Both human and macaque *CBLN2* levels increase during mid-fetal periods in parallel with key genes associated with synapse and dendrite development (**Extended Data Fig. 1**), suggesting a connection between the two events. Around the time of late infancy, both human and macaque *CBLN2* expression levels are comparable across analyzed neocortical areas, despite modest but statistically significant increased expression in PFC areas compared to non-PFC areas (Human: periods 7-10: 1.2-foldchange, periods 11-14: 1.1-foldchange; Macaque: periods 7-10: 1.25-foldchange, periods 11-14: 1.2-foldchange; **Fig. 1a**). Furthermore, the level of *CBLN2* expression is greater and the maximum expression occurs comparatively earlier in human PFC, compared to macaque PFC during mid-fetal periods (Student’s t-test, P < 0.005 at mid-fetal ages; Not significant during later fetal development to adult ages; **Fig. 1a**), suggesting precocious and transient upregulation of *CBLN2* in mid-fetal human PFC. Analysis of the spatio-temporal expression profiles of the paralogs of *CBLN2* within each species, revealed that *CBLN1* was significantly upregulated in human PFC and, to a lesser degree, in macaque PFC, compared to non-PFC areas (Human: periods 4-6: 1.59-fold change, Student’s t-test, P = 2.9e-12; Macaque: periods 4-6: 1.27-foldchange, P = 8.6e-07; **Extended Data Fig. 2**). *CBLN4* was both upregulated in the mid-fetal human PFC, compared to macaque PFC (Periods 4-6; Student’s t-test, 3.51-fold change, P = 1.5e-8), and compared to human non-PFC areas (Periods 4-6; Student’s t-test, 1.37-fold change, P = 0.002). *CBLN3* was lowly expressed in both human and macaque mid-fetal PFC (**Extended Data Fig. 2**). We also analyzed the spatio-temporal expression profiles of the genes encoding the binding partners of the CBLN family members, *NRXN1,2,3* and *GRID1,2.* While none of these genes showed a distinct PFC upregulation during mid-fetal development, all were expressed in the PFC and most showed increasing levels of expression throughout development (**Extended Data Fig. 1**).

**Figure 1.**
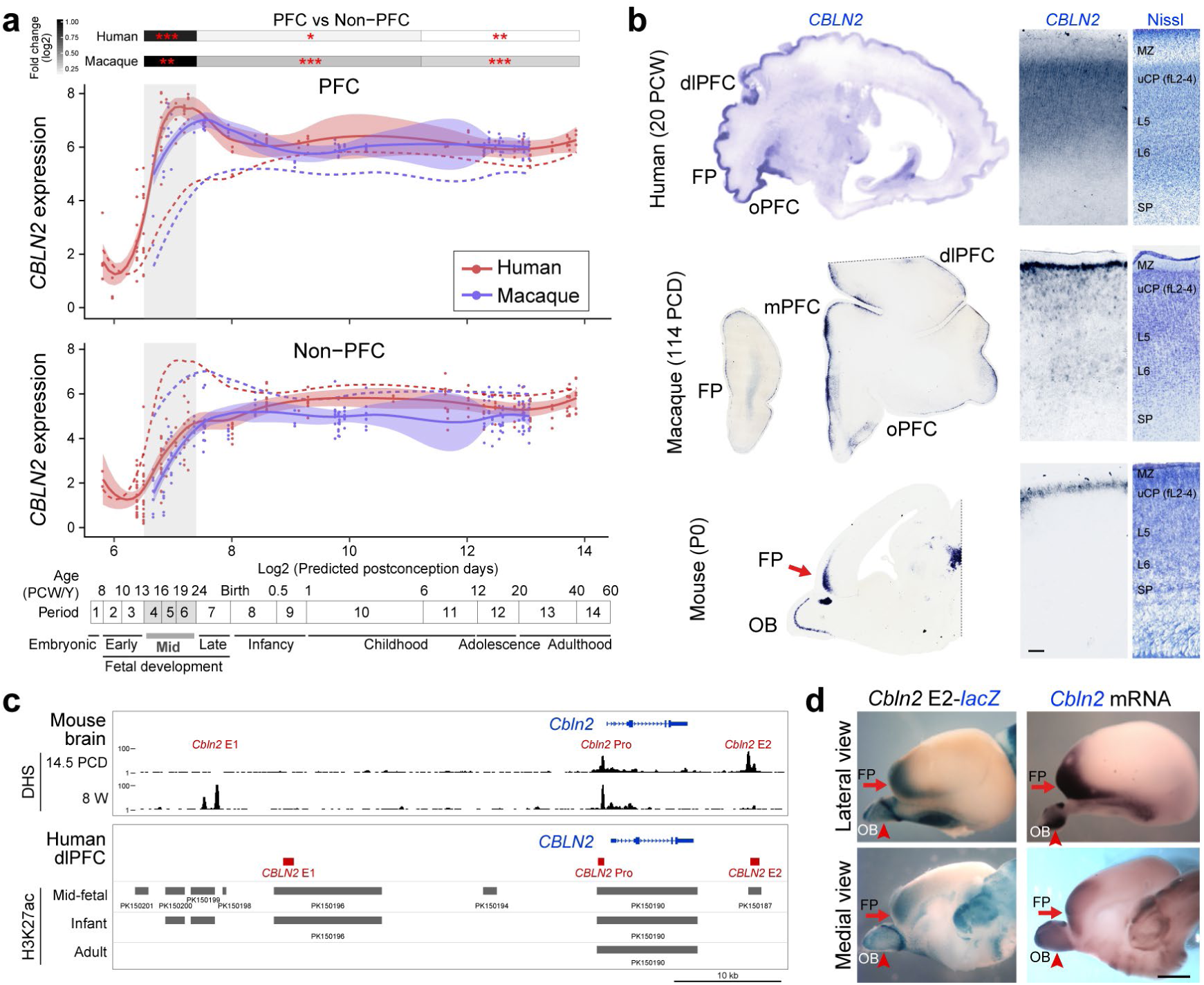
Transcriptomic and regulatory characterization of primate mid-fetal prefrontal cortex upregulation of *CBLN2*. **a**, *CBLN2* expression in the PFC and non-PFC regions of the cerebral cortex of human and macaque during development. Red and blue lines indicate human and macaque, respectively, and dotted lines represent the non-PFC *CBLN2* expression in the PFC plot and vice versa. Vertical gray box demarcates mid-fetal developmental periods. Quantification of fold change between human and macaque PFC and non-PFC *CBLN2* expression for periods 4-6, 7-10 and 11-14 (Student’s t-test; ***P *=* 2.2E-16, **P < 2E-9, *P *=* 3E-5) shown above plots. Timeline of human and macaque development and the associated periods designed by Kang et al. ^40^ shown below. Predicted ages were calculated using the *TranscriptomeAge* algorithm ^15^, which aligns our earliest macaque samples (60 PCD) with human early mid-fetal samples. **b**, *In situ* hybridization for *CBLN2* using sagittal sections of 20 PCW human and a P0 mouse and coronal sections of the PFC of 114 PCD macaque. Right panels show laminar expression of *CBLN2* in mid-fetal PFC and Nissl stain of adjacent section to show the cytoarchitecture. Sagittal human *CBLN2* image was obtained from the humanbraintranscriptome.org database ^18,40^. L, layer; MZ, marginal zone; SP, subplate; uCP, upper cortical plate. Scale bars: 500 μm **c**, DNase 1 hypersensitive sites (DHS) in whole mouse brain at 14.5 PCD and 8 postnatal week (W) within 50 kb either upstream and downstream of *Cbln2* gene. H3K27ac ChIP-seq data from human dorsolateral PFC at mid-fetal age, infancy, and adulthood (bottom panel). Putative *cis*-regulatory elements were designated as *CBLN2* E1, E2, and Pro. **d**, β-Galactosidase activity in transgenic mouse brain carrying mouse *Cbln2* E2 conjugated with *lacZ* reporter at 17 PCD. Endogenous *Cbln2* expression in age-matched brains by *in situ* hybridization is shown for comparison. Arrows and arrowheads indicate the frontal pole (FP) and olfactory bulb (OB), respectively. Scale bars: 1mm.

Previous studies have reported an anterior-posterior gradient of *CBLN2* expression in the human mid-fetal neocortex, with upregulated expression specifically in the PFC ^18,41^. These studies also showed that *CBLN2* was detected both in neurons of prospective upper (layers 2 to 4) and deeper layers (layers 5 and 6) of the human PFC, but only faintly in the more posterior mid-fetal neocortex. We confirmed this finding with close re-examination of *CBLN2* expression in the mid-fetal human PFC (21, 22 PCW; N = 2 brains), using *in situ* hybridization (**Fig. 1b**). We also identified an anterior-posterior gradient of *Cbln2* expression in macaque (114, 110 PCD; N = 2 brains) and mouse (P0; N = 4 brains) at equivalent developmental ages (**Fig. 1b**). Interestingly, in mid-fetal macaque, *Cbln2* expression was mostly observed in the upper cortical plate corresponding to the prospective upper layers with noticeably weaker expression in the deeper layers (**Fig. 1b**). In neonatal mice, *Cbln2* was restricted to the prospective upper-layers (**Fig. 1b**). Together, *CBLN2* expression displayed a progressive extension into the deeper layers of the mid-fetal PFC in humans compared to macaques, and in macaques compared to mice.

### Hominini-specific deletions in a retinoic acid-responsive *CBLN2* enhancer acting in the mid-fetal PFC

In order to understand the spatial and laminar regulatory mechanism of *CBLN2* expression in the mid-fetal PFC, we analyzed publicly available datasets on the regulatory landscape of the developing and adult mouse and human brains generated by the ENCODE and PsychENCODE projects ^20,42^. Analyzing the mouse *Cbln2* locus we identified three putative *cis-*regulatory elements marked as DNase 1 hypersensitive sites, which we designated as *Cbln2* enhancer 1 (*Cbln2* E1, 1452 bp), *Cbln2* enhancer 2 (*Cbln2* E2, 1005 bp), and *Cbln2* promoter (*Cbln2* Pro, 316 bp; **Fig. 1c, top**). The sequences of these putative *cis-*regulatory elements are moderately conserved between mouse and human (E1, 76.3%; E2, 84.3%; and Pro, 70.0%). Of note, mouse *Cbln2* E2 showed DNase 1 hypersensitivity peaks only in 14.5 postconception days (PCD) samples and *Cbln2* E1 only in 8 postnatal weeks samples (**Fig. 1c, top**). Analysis of genomic sites with differential distribution of H3K27ac, a mark of active enhancers, in the developing and adult human dlPFC indicates that *CBLN2* Pro is active at all three analyzed time points (i.e., mid-fetal, early infancy and adult ages), while *CBLN2* E1 is active during mid-fetal age and infancy, and *CBLN2* E2 is only active during mid-fetal age (**Fig 1c, bottom**).

We next assessed the activity of these putative regulatory regions using transgenic mouse lines in which a *lacZ* expression cassette was placed downstream of mouse *Cbln2* E1, *Cbln2* E2, or *Cbln2* Pro. While *Cbln2* E1-*lacZ* and *Cbln2* Pro-*lacZ* transgenic lines exhibited no detectable LacZ histochemical staining (0 of 10 founders) in the developing cortex between 17 and 18 PCD (approximately equivalent to early mid-fetal cortical development in humans), 50% of *Cbln2* E2-*lacZ* transgenics (3 of 6 founders) showed *lacZ* expression in the brain at this age, recapitulating endogenous frontal cortex-enriched *Cbln2* expression (**Fig. 1d**). These findings indicate that *Cbln2* E2 is an enhancer that drives expression in the mouse neonatal frontal cortex, including the medially located PFC areas (mPFC).

Given the presence of the *CBLN2* E2 locus in human, macaque and mouse, we hypothesized that sequence differences between the orthologs may underlie the species-specific expression pattern of *CBLN2*. Comparative genomic and phylogenetic analysis identified three separate deletions (122 bp, 20 bp and 84 bp) in the sequence of the human *CBLN2* E2 ortholog as compared to the sequence of the mouse *Cbln2* E2 (**Fig. 2a**). Further comparison across multiple vertebrates revealed that two of the three deleted sequences are jointly absent only in extant members of the Hominini clade (i.e., human, common chimpanzee, and bonobo) (**Extended Data Fig. 3a, 4**). The first two of these sequences, which we named Hominini-specific deletion 1 and 2 (HSD1 and 2, **Fig. 2a**), are highly conserved among other analyzed Haplorhini (i.e., gorilla, orangutan, gibbon, macaque, tarsier), Strepsirrhini (i.e., lemur), and mouse (**Fig. 2a, Extended Data Fig. 3, 4**). The third deletion (D3) sequence was detected only in mouse and rat of the species analyzed (**Extended Data Fig. 3b, 4**). Interestingly, we did not detect genomic regions orthologous to *Cbln2* E2 in analyzed non-placental mammals and non-mammalian chordates, although they do possess the *Cbln2* gene (**Extended Data Fig. 4**).

**Figure 2.**
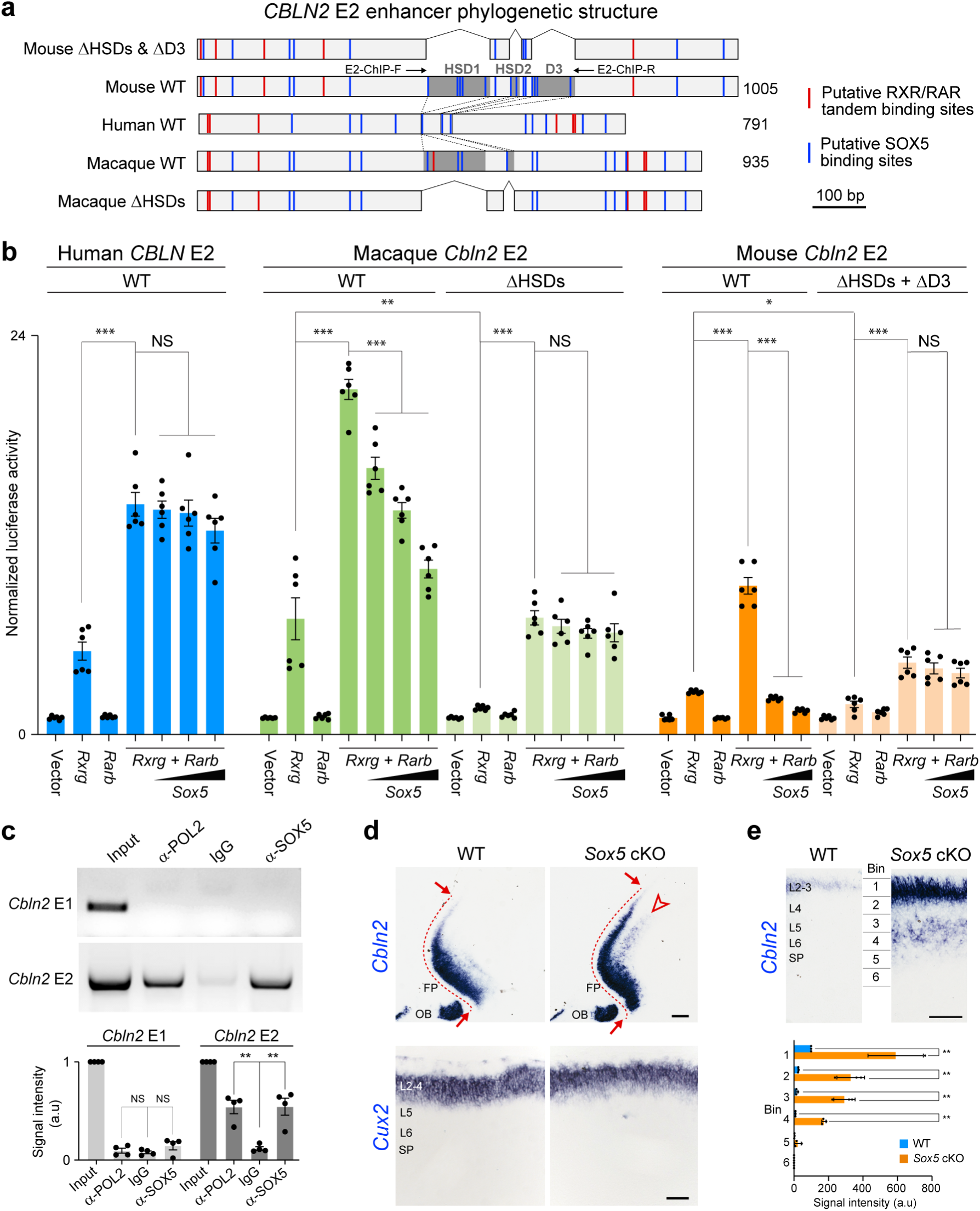
SOX5 represses *Cbln2* through Hominini-specific deletions (HSD) of regulatory sequences. **a**, Schematic representation of human, macaque and mouse *Cbln2* E2, their ΔHSDs, and ΔD3 deletion mutant constructs used for luciferase assays. Shaded dark grey areas in indicate HSD1-2 and D3 (mouse) and HSD1-2 (macaque). All HSDs and D3 are deleted from mouse and macaque *Cbln2* E2 in mouse ΔHSDs and macaque ΔHSDs. Primer set used for ChIP-PCR assays in **c** is indicated as E2-ChIP-F and R (**Supplementary Table 1**). Red and blue lines indicate putative RXR/RAR tandem and SOX5 binding sites, respectively**. b**, Luciferase assay from luciferase reporters conjugated to the wildtype human, macaque and mouse *Cbln2* E2 and mouse and macaque ΔHSD. Activation by mouse RXRG and RARB proteins was observed in all constructs, but significant suppression by mouse SOX5 protein was only seen in mouse and macaque *Cbln2* E2 reporters. Luciferase expression from mouse and macaque ΔHSDs showed no significant suppression by SOX5. Two-tailed Student’s test; ***P < 0. 0005, **P = 0. 0016, *P = 0.011; NS, not significant. Error bars: S.E.M.; N = 6 per condition. **c**, ChIP-PCR assays in P0 mouse neocortices to detect enhancer regions bound by endogenous SOX5. DNA fragments immunoprecipitated with anti-RNA polymerase II (anti-POL2, lane 2), IgG, (lane 3), and anti-SOX5 (lane 4) were detected by PCR with primers specific for the predicted *Cbln2* E1 and E2 sites. Signal intensity in each lane was quantified. Two-tailed Student’s t-test; **P < 0.005; Error bars: S.E.M.; N = 4 per condition. **d**, *Cbln2* mRNA expression in P0 *Sox5* loxP/loxP; *Emx1*-*Cre* (cKO) cortex compared to *Sox5* +/+; *Emx1*-*Cre* (WT) cortex. N = 3 per genotype. Arrows highlight posterior spread of *Cbln2* expression and arrowheads highlight ectopic deep layer expression. Upper layer marker, *Cux2*, expression was unchanged between wildtype and mutant brain. **e**, Magnification of cortical regions was divided into six bins spanning from pia to ventricular zone, and *Cbln2* signal intensity is quantified for each bin (N = 3). Two-tailed Student’s t-test; **P < 0.005. FP, frontal pole; OB, olfactory bulb; SP, subplate.

To understand transcriptional regulation of developmental *CBLN2* cortical expression, we performed transcription factor binding site analysis, which identified multiple putative RA receptor RXR/RAR tandem binding sites in *CBLN2* E2 (**Fig 2a, Extended Data Fig. 3a**). In our accompanying study ^35^, we found that *RXRG* and *RARB* are upregulated in the mid-fetal frontal lobe and that *Cbln2* expression is reduced in the neonatal frontal cortex of mice lacking both *Rxrg* and *Rarb*. Consistent with these findings, we found that overexpression of RXRG and RARB activated human, chimpanzee, gorilla, macaque, and mouse orthologs of *CBLN2* E2 in *in vitro* luciferase reporter assays more than expression of either RXRG and RARB individually (Two-tailed Student’s t-test: control vector vs. RXRG/RARB: P = 6.0e-9, 7.0e-5, 1.0e-5, 1.6e-11, 2.0e-8, for human, chimpanzee, gorilla, macaque, and mouse, respectively; **Fig. 2b; Extended Data Fig. 5a**). These observations are consistent both with our observation of higher frontal lobe expression of *Cbln2, Rxrg, and Rarb*, and previous reports showing that RXR and RAR receptors exert their effects as heterodimers in the developing and adult brain ^43,44^.

### Clade-specific suppression of *CBLN2* E2 by SOX5

In addition to RA receptors, analysis of binding sites in *Cbln2* E2 also identified multiple putative SOX5 binding sites within the enhancer, including three in HSD1 and a fourth in HSD2 (**Fig. 2a, Extended Data Fig. 3b,c**). This is significant both due to the loss of these sites in Hominini and the role of SOX5 as a primarily repressive transcription factor required for specification and development of cortical projection neurons predominantly within deep layers ^45-47^. Of note, we identified additional non-conserved SOX5 binding sites in mouse HSD1 and 2, as well as multiple SOX5 binding sites in the Muroidea specific D3 (**Fig. 2a)**. Moreover, we found that mouse *Cbln2* E2 activation by RXRG/RARB heterodimer was suppressed by SOX5 in a dose-dependent manner, whereas human and chimpanzee *CBLN2* E2 showed no significant suppression by SOX5 at the same doses (Two-tailed Student’s t-test: RXRG/RARB vs. RXRG/RARB/SOX5: P *=* 0.0004, P = 0.6, P *=* 0.7, for mouse, human, chimpanzee respectively; **Fig. 2b; Extended Data Fig. 5a**). Differential suppression of *Cbln2* E2 activation was not a consequence of sequence differences between mouse and human SOX5, as activation of mouse *Cbln2* by RXRG/RARB was suppressed by both mouse and human SOX5 (**Fig. 2b; Extended Data Fig. 5b**). Activation of macaque and gorilla *Cbln2* E2 by RXRG/RARB showed intermediate suppression of activity at the same doses of SOX5 (Two-tailed Student’s t-test: RXRG/RARB vs. RXRG/RARB/SOX5: P = 0.0004 and P = 0.01 for macaque and gorilla, respectively; **Fig. 2b; Extended Data Fig. 4a**). Deletion of HSD1, HSD2, and D3 in mouse, or HSD1 and HSD2 in macaque, abolished suppression by SOX5, suggesting that SOX5 suppression is primarily mediated by SOX5 binding sites within these HSDs and D3 (Two-tailed Student’s t-test: RXRG/RARB vs. RXRG/RARB/SOX5: P = 0.2 and 0.7 for mouse and macaque, respectively; **Fig. 2b**). To test for direct *in vivo* binding, chromatin immunoprecipitation followed by PCR (ChIP-PCR) assays were conducted using the anterior half of the neonatal mouse neocortex, which found that endogenous SOX5 bound *Cbln2* E2 but not *Cbln2* E1 (**Fig. 2c**).

To understand the functional significance of SOX5 repression of *Cbln2* E2 *in vivo*, we next examined *Cbln2* cortical expression using the *Emx1*-*Cre* transgenic mouse line engineered to achieve cortex-specific *Sox5* deletion. As predicted by luciferase assays, the absence of SOX5 protein specifically within the cortex led to ectopic expression of *Cbln2* in deeper layers of the neonatal frontal cortex (**Fig. 2d, e, Extended Data Fig. 6a**). There was also an increase and posterior extension of *Cbln2* expression in the upper layers (**Fig. 2d,e, Extended Data Fig. 6a**), likely due to previously described transient *Sox5* expression in these layers during postnatal development ^45-47^. Consistent with these results, the 5’ segment of *Cbln2* E2 lacking the two HSD sequences and D3 (*Cbln2* E2-Fr1) was not suppressed by SOX5 in luciferase assay (Two-tailed Student’s t-test: RXRG/RARB vs. RXRG/RARB/SOX5: P *=* 0.06; **Extended Data Fig. 6b,c**). Furthermore, a transgenic mouse line expressing *lacZ* under the transcriptional control of *Cbln2* E2-Fr1, showed deep layer expression of *lacZ* as compared to *Cbln2* E2 (**Extended Data Fig. 6d**). Taken together, these observations suggest SOX5 directly represses *Cbln2* expression in the deep layers of a neonatal mouse and mid-fetal macaque (and likely other non-Hominin primates) frontal cortex by binding HSD sites in *Cbln2* E2.

### Human *CBLN2* E2 increases *Cbln2* expression and synaptogenesis in mouse PFC

To test whether human *CBLN2* E2 is sufficient to drive a human-like pattern of *Cbln2* expression in the mouse neonatal frontal cortex, we generated a mouse line in which the *Cbln2* E2 enhancer was replaced *in situ* (i.e., in the orthologous position) with the corresponding human *CBLN2* E2 enhancer using the CRISPR-Cas9 genome editing technique (**Extended Data Fig. 7**). The engineered mice carrying homozygous humanized *Cbln2* E2 (h*Cbln2* E2) in both alleles (*Homo sapiens*;*Homo sapiens, Hs;Hs*) were viable and fertile, and the expression of the cortical upper layer specific marker, *Cux2*, and SOX5 were comparable to homozygous WT *Cbln2* E2 (*Mus musculus;Mus musculus, Mm;Mm*) mice (**Fig. 3c)**, indicating that genetically humanized h*Cbln2* E2grossly functions as the mouse ortholog. Neocortical *Cbln2* expression at P0 was increased by 21.7% in h*Cbln2* E2 (*Hs;Hs*) cortex, as compared to WT mouse *Cbln2* E2 (*Mm;Mm*) by quantitative RT-PCR (Two-tailed Student’s t-test: WT *Cbln2* E2 vs. h*Cbln2* E2: P *=* 0.007; **Extended Data Fig. 8**). Analysis of expression by *in situ* hybridization showed that *Cbln2* was also transiently upregulated in both upper and deeper layers of the h*Cbln2* E2 PFC at 18 PCD and P1, but not at 16 PCD (**Fig. 3a,b; Extended Data Fig. 9**). After P1, *Cbln2* expression extended into the deeper layers of PFC, coinciding with the down-regulation of SOX5 protein expression (**Extended Data Fig. 10**) in both h*Cbln2* E2and WT *Cbln2* E2.

**Figure 3.**
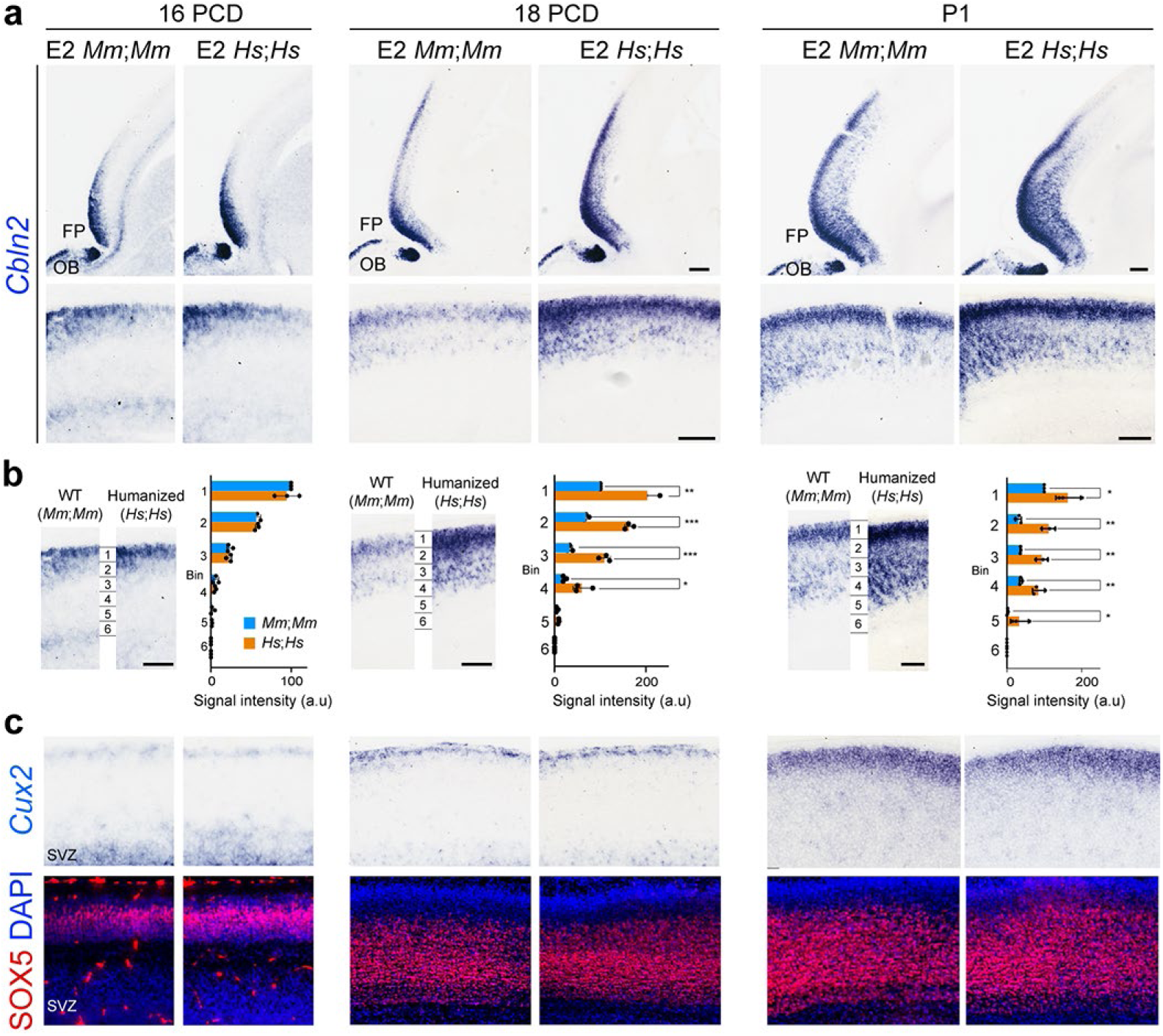
Humanized *Cbln2* E2 drives upregulation and ectopic laminar *Cbln2* expression in the neonatal anterior/frontal cortex. **a**, *Cbln2* expression in the humanized *Cbln2* E2 knock-in mouse (h*Cbln2* E2, *Hs;Hs*) brain at 16 PCD, 18 PCD and P1 compared to WT mouse (WT *Cbln2* E2, *Mm;Mm*) with higher magnification panels below. **b**, Cortex was divided into six equal bins spanning from pia to ventricular zone, and *Cbln2* signal intensity was quantified for each bin and compared between WT *Cbln2* E2 (*Mm;Mm*) and h*Cbln2* E2 (*Hs;Hs*). Two-tailed Student’s t-test; *P < 0.05, **P < 0.005, ***P < 0.0005; N = 3 per genotype. Upregulated *Cbln2* expression in both upper and deeper layers is observed in h*Cbln2* E2 brains from 18 PCD and P1, but not in 16 PCD. **c**, Expression of the upper layer marker, *Cux2*, and SOX5 in adjacent tissue sections were detected by *in situ* hybridization and immunostaining, respectively. Scale bars: 100 μm.

Members of the CBLN family have been reported to regulate synapse development and function ^21-27^. Therefore, we hypothesized that a transient increase in *Cbln2* expression in the h*Cbln2* E2 PFC will lead to increased synapse number. Although *Cbln2* is primarily expressed by glutamatergic excitatory projection neurons in the mouse mPFC ^27^, it can act as an organizer for both excitatory and inhibitory synapses ^23-27^. Thus, we quantified both excitatory and inhibitory synaptic density in the mPFC, primary somatosensory area (SSp), and primary visual (VISp) area using immunostaining for postsynaptic proteins PSD-95/DLG4 and gephyrin, respectively. In E2 *Hs;Hs* mice, we observed an increase in the density of PSD-95-immunopositive excitatory postsynaptic puncta in both upper (39.5%, BCL11B-immunonegative; two-tailed Student’s t-test: WT *Cbln2* E2 vs. h*Cbln2* E2: P = 0.0004; N = 5) and deep layers (47.9%, BCL11B-immunopositive; two-tailed Student’s t-test: WT *Cbln2* E2 vs. h*Cbln2* E2: P = 0.0001; N = 5) in the mPFC but not in the SSp and VISp, where *Cbln2* expression is not altered, at P1 (**Fig. 4a,b**). A similar increase in synaptic density in the mPFC was identified in the *Sox5* conditional KO mice (58% for upper layers; two-tailed Student’s t-test: WT *Cbln2* E2 vs. h*Cbln2* E2: P = 2E-5, and 40% for deep layers; two-tailed Student’s t-test: WT *Cbln2* E2 vs. h*Cbln2* E2: P = 3E-5; N = 3, **Fig. 4b**). Similarly, we also observed an increase in the density of gephyrin-immunopositive inhibitory postsynaptic puncta number in both upper (24.2%, BCL11B-immunonegative; Two-tailed Student’s t-test: WT *Cbln2* E2 vs. h*Cbln2* E2: P = 0.0002; N = 3) and deep layers (37.9%, BCL11B-immunopositive; two-tailed Student’s t-test: WT *Cbln2* E2 vs. h*Cbln2* E2: P = 5E-5; N = 3) at P0 in the h*Cbln2* E2PFC but not in the SSp and VISp cortex (**Fig. 4c,d**). In the *Sox5* conditional KO mice, anti-gephyrin immunostaining revealed a similar increase in the density of inhibitory synapses (27.3% for upper layers, two-tailed Student’s t-test: WT *Cbln2* E2 vs. h*Cbln2* E2: P = 0.0002; 39.4% for deep layers, two-tailed Student’s t-test: WT *Cbln2* E2 vs. h*Cbln2* E2: P = 0.0003; N = 3, **Fig. 4d**). Interestingly, at P60, an increase in the density of PSD-95-immunopositive synaptic puncta was observed only in deep layers of humanized *CBLN2* E2 mice (Two-tailed Student’s t-test: WT *Cbln2* E2 vs. h*Cbln2* E2: P = 0.02; 21.4%; N = 3), but not in upper layers (**Fig. 4b**), and no significant changes were observed in the density of gephyrin-immunopositive synaptic puncta at this age (**Fig. 4d**).

**Figure 4.**
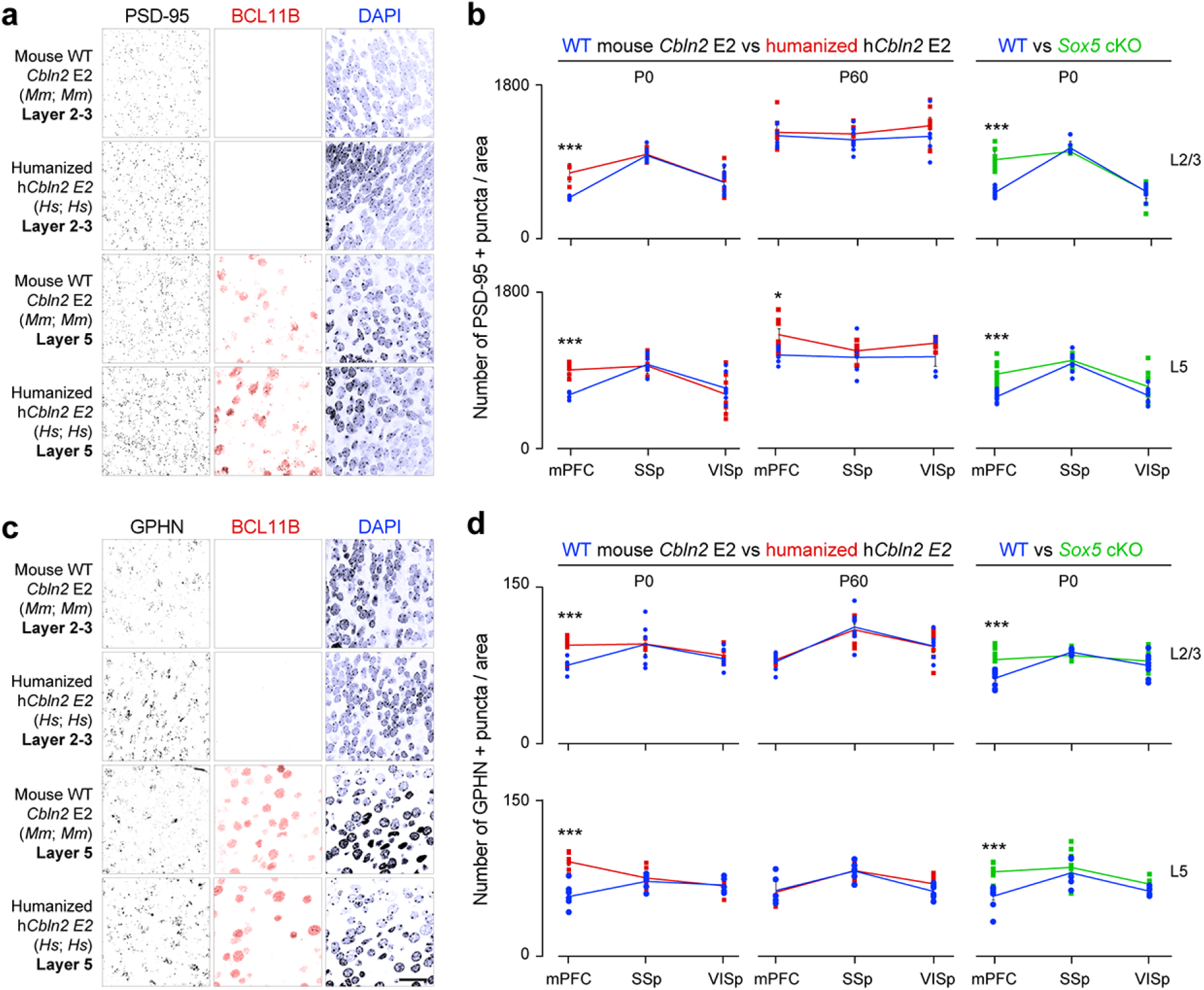
Increased density of prefrontal excitatory and inhibitory postsynaptic puncta in humanized *CBLN2* enhancer mice. Representative images of PFC layer 2/3 (BCL11B-immunonegative) and layer 5 (BCL11B-immunopositive) from WT *Cbln2* E2 (*Mm;Mm*) and h*Cbln2* E2 (*Hs;Hs*) with immunostaining with anti-PSD-95/DLG4 (**a**) and anti-Gephyrin (GPHN; **c**) antibodies at P0. Quantification of density of excitatory postsynaptic puncta marked by PSD-95/Dlg4 (**b**), and inhibitory postsynaptic puncta marked by GPHN (**d**) from layer 2/3 (L2/3) and layer 5 (L5) of the mPFC, primary somatosensory area (SSp), and primary visual area (VISp) at P0 and P60 WT *Cbln2* E2 (blue), h*Cbln2* E2(orange), or *Sox5* conditional KO mice (green) at P0. Two-tailed Student’s test; *P = 0.018; ***P < 0.0005; Error bars: S.E.M.; N = 5 per genotype for P0 brain; N = 3 per genotype for P60 brain. P60 brains were all from females. Scale bar: 25 μm.

## Conclusion

In this study, we describe a plausible molecular mechanism and underlying genetic basis for the anterior-posterior gradient of cortical synaptogenesis and disproportionate increase in the number of synapses within the human PFC. Using a spatio-temporal cortical transcriptomic database and multi-species analysis, we identified a transient upregulation of synaptic organizer *CBLN2* in the human PFC during mid-fetal development, which coincides with the initiation of synaptogenesis and cortical circuitry formation ^48^. The mid-fetal upregulation of *CBLN2* expression is greater in amplitude and occurs earlier in humans compared to macaque, identifying a putative heterochronic expression of *CBLN2* in humans. Furthermore, we identified an expansion of *CBLN2* into the deeper layers of the mid-fetal human PFC compared to macaque and mouse PFC, which is the result of Hominini-specific deletions in the PFC-active *CBLN2* enhancer containing SOX5 binding sites. Mutations associated with SOX5 have been linked to multiple neuropsychiatric and neurodevelopmental disorders, including ASD, and have recently been shown to have broad cross-disorder association ^49-52^.

Interestingly, *Cbln2* expression extends into deep layers of postnatal mice, thus suggesting a period of synaptogenesis in these layers and, through comparison with human data presented in this study, identifying periods where synaptogenesis may be extended specifically in Hominini in those layers of the PFC. Accordingly, the difference in expression level of *CBLN2* between PFC and non-PFC regions is lost around birth, prior to the peak of cortical synaptogenesis in both humans and macaques ^33,34^, supporting the concept of regional heterochronic initiation of synaptogenesis, specifically in the PFC ^32^. This transient developmental upregulation of *CBLN2*, when modelled in mouse using either *in situ* humanization of the mouse *CBLN2* E2 or the elimination of SOX5-dependent repression predicted to occur in human development, led to an increase in the density and likely the total number of excitatory and inhibitory synapses in both upper and deep layers of the mPFC in early postnatal development, and an increase of only deep layer excitatory synapses in the adult. Furthermore, in the adult cortex, the humanization of the mouse *CBLN2* E2 creates an anterior to posterior gradient of excitatory synapses, not present in WT at the analyzed ages (**Fig. 4b**). Thus, transient mid-fetal Hominini-specific PFC upregulation of *CBLN2* may underlie increased synaptic connectivity in the human PFC. In our accompanying paper ^35^, we identified that loss of RA signaling in the developing mouse mPFC leads to both reduced expression of *Cbln2*, as well as of its functionally related paralog *Cbln1*, and reduced density of excitatory synapses. Regulation of *Cbln2* E2 by RA receptors, RXRG and RARB, provides a direct link between these two finding, and may be part of a more complex gene regulatory network regulating development and adaptive changes in PFC connectivity.

While CBLN2 has been implicated in Tourette’s syndrome and OCD ^37^ and mouse CBLN1 is regulated by ASD- and Angelman syndrome-linked UBE3A ^53^, they both serve as ligands for multiple NRXNs and GRID1, which are ASD- and schizophrenia-associated proteins, respectively ^36-39,54^. Multiple lines of evidence have shown that ASD and schizophrenia risk genes, converge in time, brain region, cell type and cell compartment with strong evidence for convergence in synapses ^38,39^ as well as glutamatergic projection neurons and frontal cortex during human mid-fetal development ^36,55-58^. Consistent with these findings, alterations in the levels of some synaptic proteins, the number of synapses, and the amount of connectivity within frontal lobe have been described as the possible underlying pathophysiology of both ASD and schizophrenia ^59,60^ with dysfunction in the NXRN-GRID trans-synaptic complex, which includes CBLNs, being a putative mechanism and target for interventions.

## Methods

### Analysis of human and macaque transcriptomic data

RNA-seq data from developing human and macaque brains was obtained from Zhu et al. ^15^. Predicted ages for macaque samples were calculated via the *TranscriptomeAge* algorithm described in Zhu et al. ^15^. To perform statistical comparisons, samples from various developmental periods were grouped (periods 4-6, 7-10, and 11-14) and a two-tailed Student’s t-test was used to compare gene expression levels between brain regions and species. Sequencing data were deposited at http://psychencode.org and NCBI dbGAP Accession phs000755.v2.p1.

### Animals

All experiments using animals were performed in accordance with protocols approved by the Yale University Institutional Animal Care and Use Committee (IACUC). The day on which a vaginal plug was observed was designated as 0.5 PCD. Mice carrying the floxed *Sox5* allele were a kind gift from Véronique Lefebvre ^61^.

### Plasmid construction

For construction of expression vectors used for luciferase assays, full-length cDNAs (Human *SOX5*, clone ID 30343519; mouse *Rarb*, clone ID 30608242; mouse *Rxrg*, clone ID 5707723; all purchased from GE Healthcare) were inserted into pCAGIG vector (Addgene plasmid # 11159). For luciferase reporter plasmids, human, chimpanzee, gorilla, macaque, and mouse *Cbln2* E2 fragments were PCR-amplified from genomic DNA of individual species and inserted into pGL4.24 vector (Promega). The sequences of the PCR primers and synthetic oligonucleotides are listed in **Supplementary Table 1**.

### Generation of transgenic mouse

*Cbln2* E2 and its related fragments were PCR-amplified from mouse genomic DNA and ligated into pBgn-*lacZ* ^62^. Vector was linearized with BglII and KpnI and purified by gel separation followed by phenol/chloroform extraction. Transgene fragment was diluted into microinjection buffer (5 mM Tris-HCl, 0.1 mM EDTA), and injected into the pronucleus of FVB mouse fertilized eggs. Transgenic mice were screened for the presence of transgenes by PCR using the *lacZ* primer set listed in **Supplementary Table 1**.

### Generation of humanized *Cbln2* E2 mouse

The overall strategy for the generation of humanized *Cbln2* E2 (h*Cbln2* E2) follows a previously described protocol ^63^. The targeting vector carrying h*Cbln2* E2 mouse was constructed as follows: mouse genomic DNA fragments of 1601 bp (chr18: 86721709-86723309, GRCm38/mm10) and 1909 bp (chr18: 86724521-86726429, GRCm38/mm10) flanking the region containing *Cbln2* E2 (chr18: 86723310-86724520, GRCm38/mm10) were PCR-amplified as left and right arm, respectively, using mouse genomic DNA as template. An 852 bp fragment containing human *CBLN2* E2 (chr18: 72530473-72531324, GRCh38/hg38) was PCR-amplified using human genomic DNA as a template. These three DNA fragments were ligated into *XhoI, HindIII* and *ClaI* sites of the PL451 vector ^64^. For the construction of the templates of guidance RNA, two sets of top and bottom strand oligomers (see **Supplementary Table 1**) directing the double strand break at mouse left and right arm were annealed and ligated into *BbsI* site of pX330 vector ^65^. After amplification of insert with T7-tagged primers (see **Supplementary Table 1**), guidance RNAs were synthesized by T7 RNA polymerase. The coding region of Caspase9 was PCR-amplified using pX330 as a template and inserted into the pSP64 Poly(A) vector (Promega). Vector were digested and linearized with *EcoRI*. Caspase mRNA was synthesized by SP6 RNA polymerase. Guidance RNAs and Cas9 mRNA were purified by MEGAclear Transcription Clean-Up Kit (Ambion). The targeting vector, Cas9 mRNA, and two guidance RNAs were mixed at a concentration of (10 ng; 100 ng; 100 ng; 200 ng μl^-1^) in the microinjection buffer (5mMTris-HCl pH7.5; 0.1M EDTA) and injected into the pronuclei of fertilized eggs from the B6SJLF1/J mouse strain. The first generation (F0) mice with recombined allele were identified by long-distance PCR with a primer set (mP3/mP4) designed outside of targeting vector (**Supplementary Table 1, Extended Data Fig. 7a**), and confirmed by sequencing. The germ line transmission in F1 generation was confirmed by nested PCR using the primer set of mP3/mP4, followed by hP1/P2 (**Extended Data Fig. 7b,c**), to exclude the possibility of detecting targeting vector randomly integrated into the genomic DNA. Mice in the following generation were genotyped by PCR with mP1/mP2 and hP1/P2 as indicated in **Extended Data Fig. 7**.

### *In situ* hybridization

Whole-mount and section *in situ* hybridization were performed as described previously ^66^. Antisense digoxigenin (DIG)-labeled RNA probes were synthesized using DIG or Fluorescein RNA Labeling Mix (Roche). Human and mouse *Cbln2* cDNA, and mouse *Cux2* cDNA were obtained from GE Healthcare (Clone ID 5727802, 6412317, and 30532644, respectively). Macaque *Cbln2* cDNA was synthesized using total RNA from adult macaque dlPFC region. The PCR fragment was ligated into pCRII vector (Invitrogen). Tissue sections were obtained from 21 and 22 PCW de-identified postmortem human brains, 110 and 114 PCD postmortem macaque brains, and mouse postmortem brains of various developmental ages. *In situ* hybridization experiments were repeated using these two sets.

### Immunohistochemistry

Postmortem brains were dissected and fixed with 4% paraformaldehyde overnight at 4°C, followed by embedding in OCT. Brains were sectioned at 15-20 µm by cryostat (Leica CM3050S) after they were frozen. For antigen retrieval, sections were treated with R-Buffer AG (Electron Microscopy Sciences) for 20 min at 120 °C. The density of excitatory and inhibitory postsynaptic puncta were quantified using immunostaining of PSD-95 (also known as Dlg4) ^67^ and Gephyrin (GPHN) ^68^, respectively. The sources of primary antibodies were anti-PSD-95/DLG4 (1:500; Invitrogen), anti-gephyrin (1:500; Synaptic Systems), anti-SOX5 (1:500; Abcam), anti-BCL11B (1:1000; Abcam). Secondary antibodies: Alexa Fluor 488-, or 594-conjugated AffiniPure Donkey anti-Rabbit IgG (Jackson ImmunoResearch). For all microscopic analysis, LSM META (Zeiss), and LSM software ZEN were used.

### Quantification of post-synaptic puncta

For each cortical area, using both the 488 nm channel to detect PSD-95 (also known as DLG4) or gephyrin (GPHN) and the 594 nm channel to detect BCL11B, seven serial optical sections at 0.8 µm intervals over a total depth of 5 µm were imaged and the 2^nd^, 4^th^, and 6^th^ images were eliminated from further analysis to avoid overlap in counting ^69^. Area of each image is 0.079 mm^2^. The number of PSD-95 or Gephyrin immunopositive puncta on each image was automatically counted using ImageJ. BCL11B-immunonegative cells were considered as layer 2-4, and BCL11B-immunopositive nuclei were considered as layer 5. At least three sections from each animal were selected for counting, and at least 3 animals for each genotype were used.

### Processing, analysis, and image visualization

To allow robust visualization and analysis, images depicting immunohistochemistry using antibody against PSD95 and gephyrin have been inverted and/or pseudo colored, as in Fig. 4.

### ChIP-sequencing data analysis

H3K27ac ChIP-seq from the developing human brain was obtained and peaks were called by Li et al. ^20^ that were converted to the mm10 genome build using LiftOver ^70^ and visualized using the Integrative Genomics Viewer ^71^. DNase 1 hypersensitivity annotation was obtained and visualized via the UCSC Genome Browser ^72,73^.

### Luciferase assays

Neuro-2a cells were transfected using Lipofectamine 2000 (Invitrogen) with either mouse or human pCAGIG-*Sox5, Rxrg, Rarb*, or empty pCAGIG, together with one of the pGL4 (Promega) luciferase vectors generated with enhancer sequences as described above. *Renilla* luciferase plasmid (pRL, Promega) was co-transfected to control for transfection efficiency. The luciferase assays were performed 48 h after transfection using the dual-luciferase kit (Promega) according to the manufacturer’s instructions. Luciferase activity were measured and quantified by GloMax®-Multi Detection System (Promega).

### Chromatin immunoprecipitation (ChIP)

The anterior half of cortices from P0 mice were dissociated then cross-linked with 1% formaldehyde and processed using EZ-ChIP kit (Millipore). Chromatin fragments bound by endogenous mouse SOX5 were pulled down by anti-SOX5 (Abcam: ab94396), anti-POL2 antibody, or random IgG control then detected by PCR using E2-ChIP-F and E2-ChIP-R or E1-ChIP-F, and E1-ChIP-R (see **Supplementary Table 1**).

### Quantitative reverse transcription-PCR

Total RNA was isolated using Trizol (ThermoFisher) from freshly dissected neocortices after removal of the olfactory bulb, hippocampus, and striatum. cDNAs were prepared using SuperScript II (Invitrogen) from more than three independent WT littermates and humanized *Cbln2* E2 brains. Quantitative reverse transcription-PCR was performed as described previously ^74^. At least three biological replicates per transcript were used for every reaction. The copy number of transcripts was normalized against the house keeping TATA-binding protein (TBP) transcript level. For *Cbln2* and *Tbp* primer sets, correlation (R2) was higher than 0.98, and the slope was −3.1 to −3.6 in each standard curve. Primers were designed in a single exon and are listed in **Supplementary Table 1**.

## Acknowledgements

We thank Suxia Bai, Timothy Nottoli and Xiaojun Xing for technical help with generating transgenic mice and genetically humanized mice; Andre Sousa for providing tissue; Belen Lorente-Galdos and Gabriel Santpere for helping with data analysis; Alvaro Duque for using equipment from MacBrainResource (MH113257); Véronique Lefebvre for providing floxed *Sox5* mice; and the members of the Sestan laboratory for comments. This work was supported by the National Institutes of Health (MH106874, MH106934, MH110926, MH116488), and the Simons Foundation (N.S.). Additional support was provided by the National Science Foundation Graduate Research Fellowship Program (S.K.M.), the Kavli Foundation and the James S. McDonnell Foundation (N.S.).

## Author Contributions

M.S., K.P., and N.S. designed research; M.S. and K.P. performed overall experiments, analyzed the data; S.K.M. analyzed RNA-seq, ChIP-seq and genomic sequence data; M.S. generated transgenic and knock-in mice; N.S. conceived the study; M.S., K.P., and N.S. wrote the manuscript. All authors discussed the results and implications and commented on the manuscript at all stages.

---

**Extended Data Figure 1.**
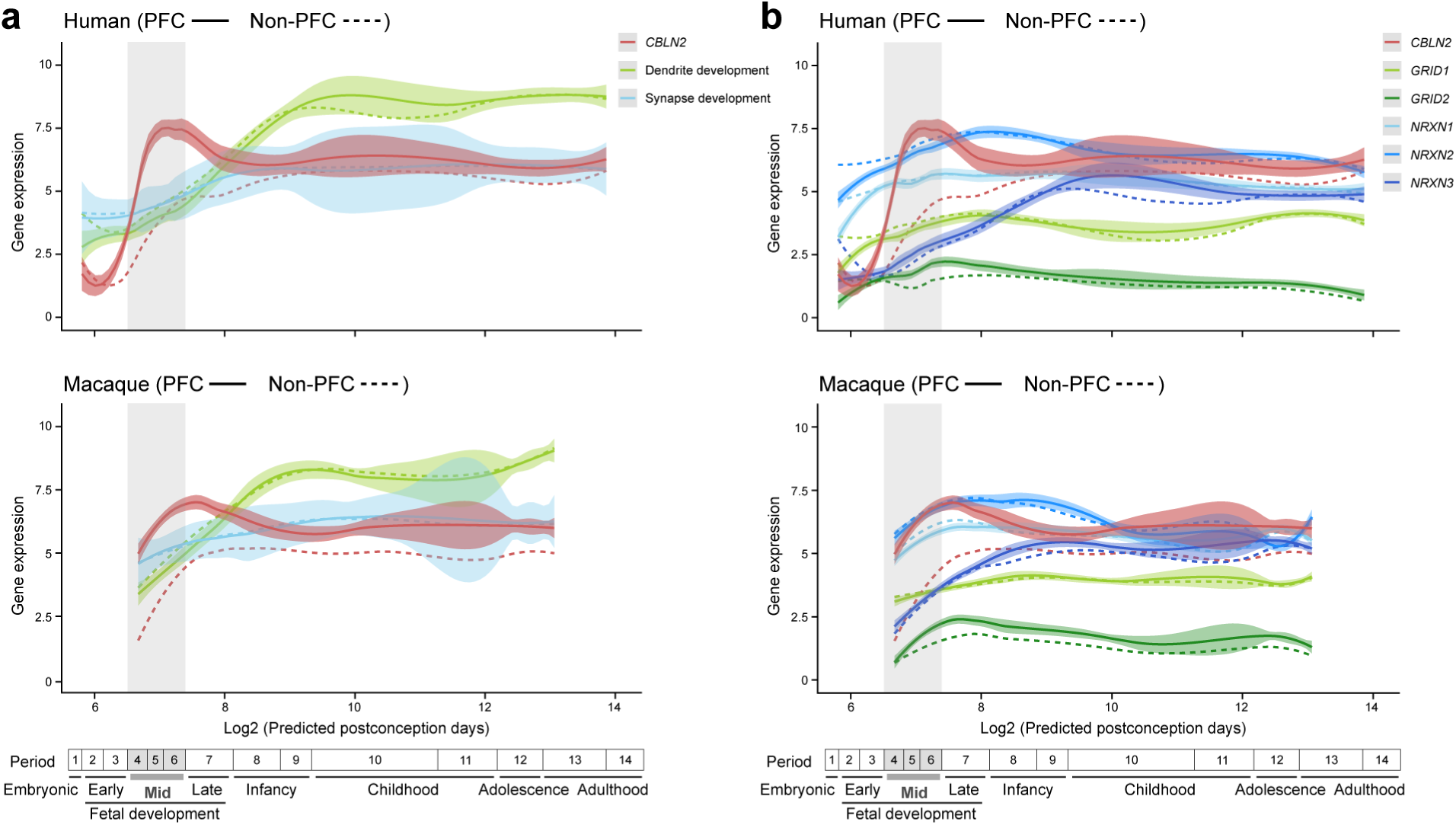
Expression profile of genes related to synapse and dendrite development, and CBLN2 binding partners in macaque and human. **a**, Developmental trajectory of *CBLN2* expression compared to the expression of key genes related to dendrite development (i.e., *MAP1A* and *CAMK2A*) and synapse development genes (i.e., *SYP, SYPL1, SYPL2* and *SYN1*) in the PFC (solid line) and non-PFC areas (dashed line) of the human (top) and macaque (bottom) cerebral cortex. The lists of genes related to synapse and dendrite development were previously complied and analyzed for their expression trajectories by Kang et al. ^40^. **b**, Developmental trajectory of *CBLN2* expression compared to the expression of genes encoding CBLN2 binding partners (i.e., *GRID1, GRID2, NRXN1, NRXN2*, and *NRXN3*) in the PFC (solid line) and non-PFC (dashed line) areas in humans (top) and macaques (bottom). Predicted ages were calculated using the *TranscriptomeAge* algorithm ^15^, which aligns our earliest macaque samples (60 PCD) with human early mid-fetal samples. Gene expression values are represented as log2(RPKM+1). Vertical grey box highlights the mid-fetal developmental periods. Timeline of human and macaque development designed by Kang et al. ^40^ shown below.

**Extended Data Figure 2.**
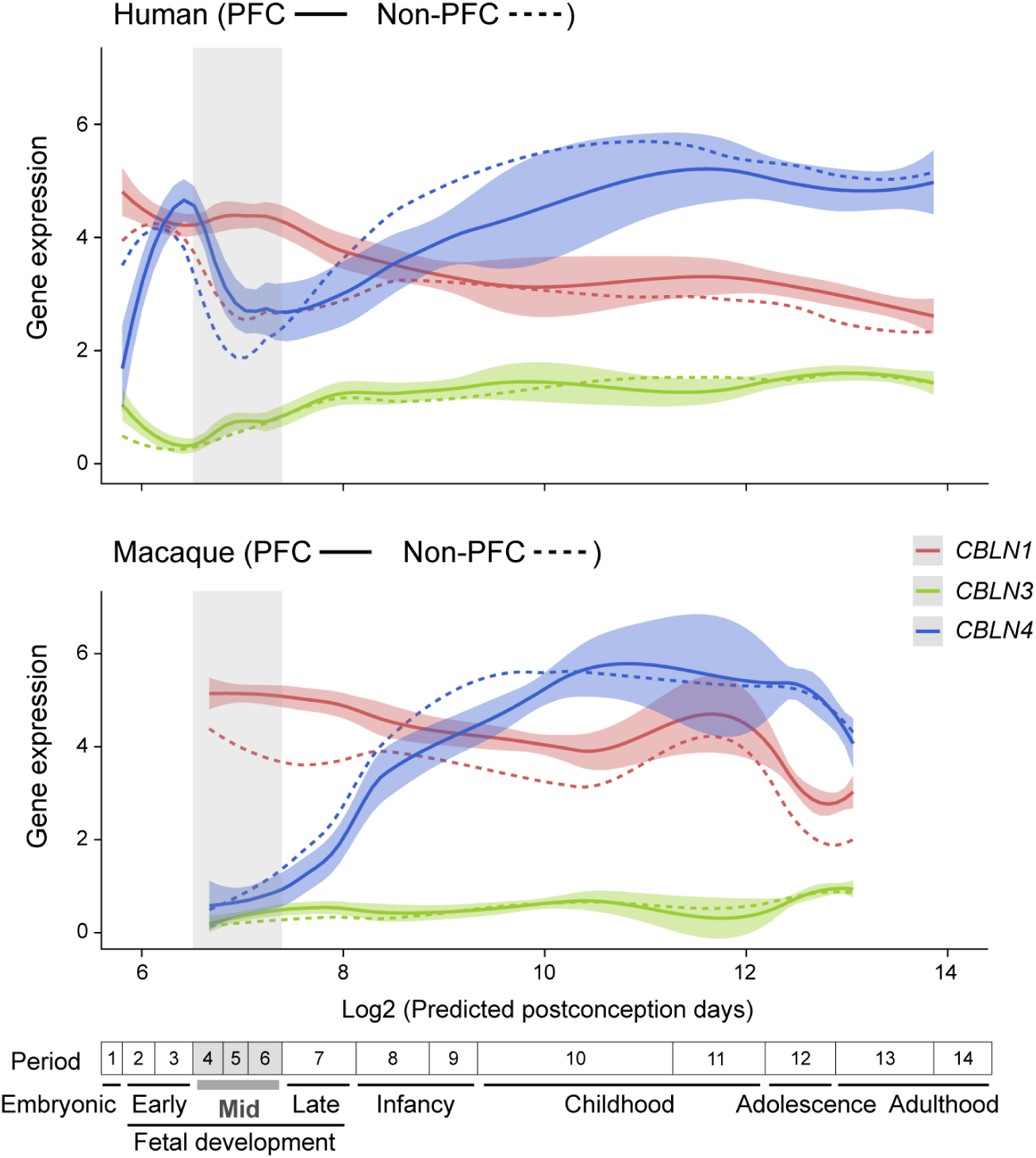
*CBLN1, 3*, and *4* expression in the developing human and macaque cerebral neocortex. *CBLN1, 3*, and *4* expression in the PFC and non-PFC areas of the developing cerebral cortex. Gray box demarcates mid-fetal development. Timeline of human and macaque development and the associated periods designed by Kang et al. ^40^ is provided in the bottom. *CBLN1* is significantly upregulated in human PFC and, to a lesser degree, in macaque PFC, compared to non-PFC areas (Human: periods 4-6: 1.59-fold change, Student’s t-test, P = 2.9e-12; Macaque: periods 4-6: 1.27-fold change, P = 8.6e-07; **Extended Data Fig. 2**). *CBLN4* was both upregulated in the mid-fetal human PFC, compared to macaque PFC (Periods 4-6; Student’s t-test, 3.51-fold change, P = 1.5e-8), and compared to human non-PFC areas (Periods 4-6; Student’s t-test, 1.37-fold change, P = 0.002). *CBLN3* was lowly expressed in both human and macaque mid-fetal PFC. Abbreviations were as described in Fig. 1 legend.

**Extended Data Figure 3.**
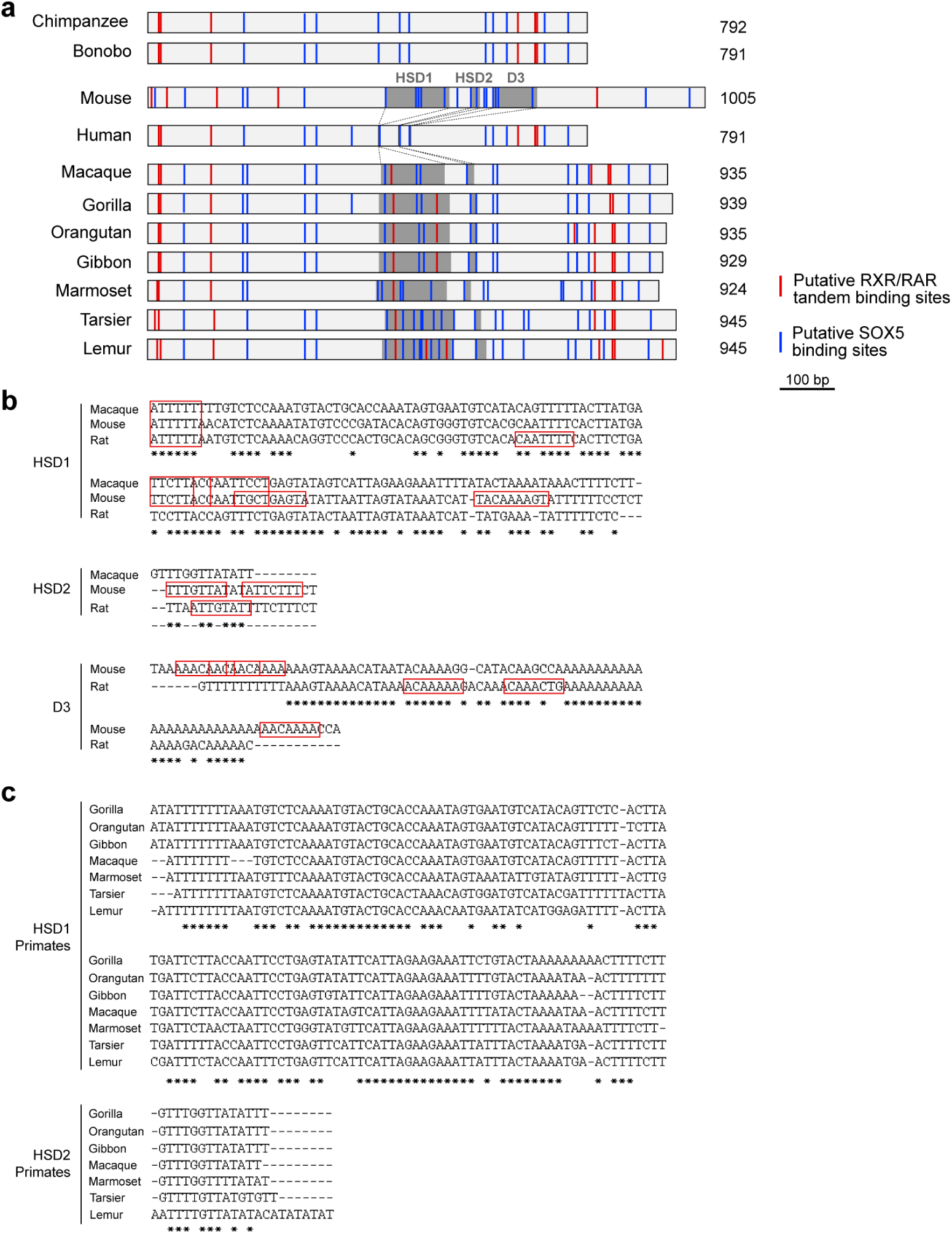
Comparative analysis of *CBLN2* E2 deletions across mammals. **a**, Schematic representation of *CBLN2* E2 from mouse and primate species including apes (human, common chimpanzee, bonobo, gorilla, orangutan, and gibbon), old world monkey (Rhesus macaque), new world monkey (marmoset), and prosimian (tarsier and lemur). HSD1, 2 and D3 are shaded. Putative RXR/RAR tandem binding sites indicated as red lines and putative SOX5-binding sites as blue lines. **b**, Sequence alignments of HSD1 and 2, and D3 from macaque, mouse, and rat. Putative SOX5 binding sites are shown in red boxes. **c**, Sequence alignments of HSD1 and HSD2 from primates shown in (**a**).

**Extended Data Figure 4.**
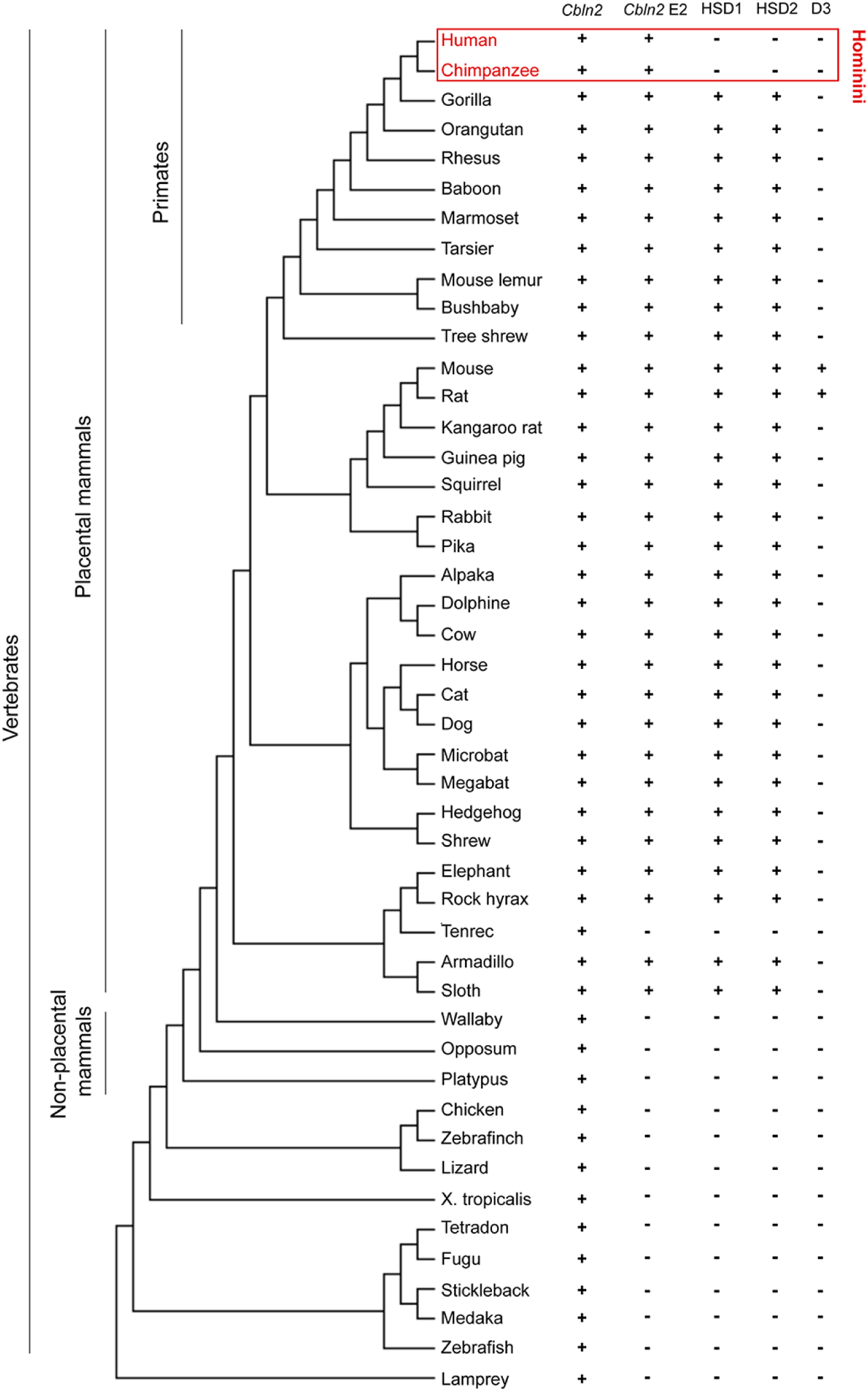
Conservation of *CBLN2* E2 and deleted regions across species. Phylogenetic tree of selected chordates including placental and non-placental mammals with information about presence of the *CBLN2/Cbln2* gene and *CBLN2* E2/Cbln2 E2. The last three columns describe the presence of HSD 1,2, and D3. Deletion of HSD1/2 is specific to humans and chimpanzees (i.e., common chimpanzee and pygmy chimpanzee or bonobo) grouped as Hominini. The mouse E2 sequence was searched in each most updated animal genome browser at UCSC genome browser (as of December 22, 2019). E2 conservation criteria are: 1) identity over 80%; and 2) alignment length over 600 bp as compared with mouse E2 (1005 bp).

**Extended Data Figure 5.**
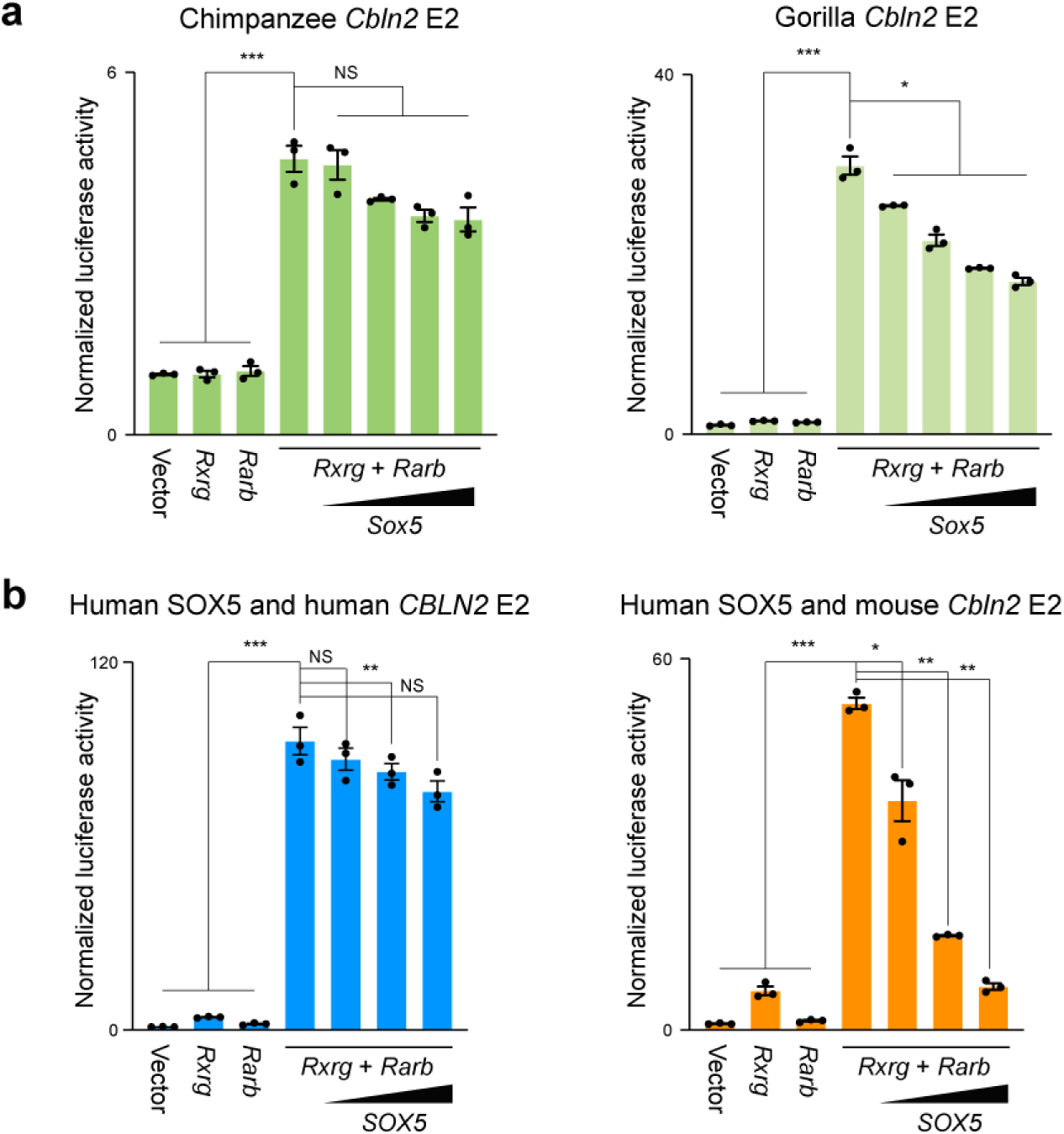
SOX5 suppresses RXRG/RARB responsive human, chimpanzee, gorilla and mouse *Cbln2* E2 enhancer. **a**, Luciferase assay shows chimpanzee *Cbln2* E2 is not suppressed by SOX5, but gorilla *Cbln2* E2, which possesses HSDs is suppressed by SOX5. Two-tailed Student’s test; *P < 0.05; **P < 0.005; ***P < 0.0005; NS, not significant. Error bars; S.E.M.; N = 3 per condition. **b**, Overexpression of human *SOX5* exerts a similar effect to mouse *Sox5* on human and mouse *Cbln2* E2 reporters; only minimal suppression of human *CBLN2* E2, but strong suppression of mouse *Cbln2* E2. Two-tailed Student’s test; *P < 0.05; **P < 0.005; ***P < 0.0005; NS, not significant; Error bars, S.E.M.; N = 3 per condition.

**Extended Data Figure 6.**
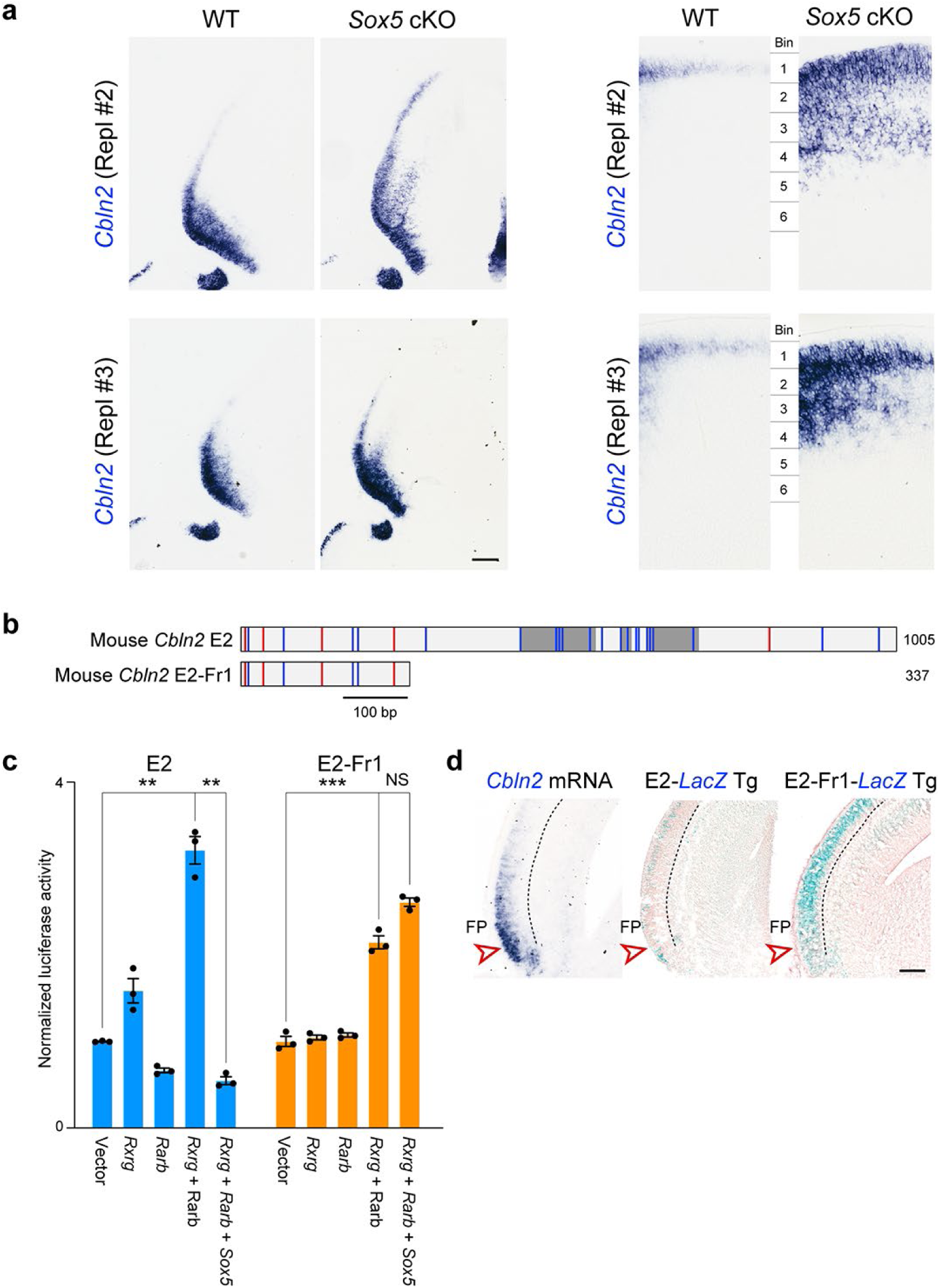
*Cbln2* expression is upregulated in *Sox5* conditional knockouts and *Cbln2* E2 5’ fragment transgenic lines. **a**, *Cbln2* expression is upregulated in *Sox5* conditional knockout brain at P0. Additional two replicates (Repl #2 and 3) not shown in Figure 2d are shown. Scale bar: 200 μm. **b**, Constructs used for luciferase assay and generation of transgenic animals. **c**, Luciferase assay for *Cbln2* E2 and *Cbln2* E2-Fr1. *Cbln2* E2-Fr1 reporter was activated by RXRG and RARB, but not suppressed by SOX5. Two-tailed Student’s test; **P < 0.005, ***P = 0.00016; NS, not significant; Error bars: S.E.M.; N = 3 per condition. **d**, Transgenic mouse brain at 17 PCD carrying *Cbln2* E2 (N = 3) or *Cbln2* E2 Fr1-*lacZ* reporters (N = 3). *Cbln2* expression in the PFC is indicated by arrowheads. Endogenous *Cbln2* expression is also shown for comparison (N = 4). Scale bar: 200 μm.

**Extended Data Figure 7.**
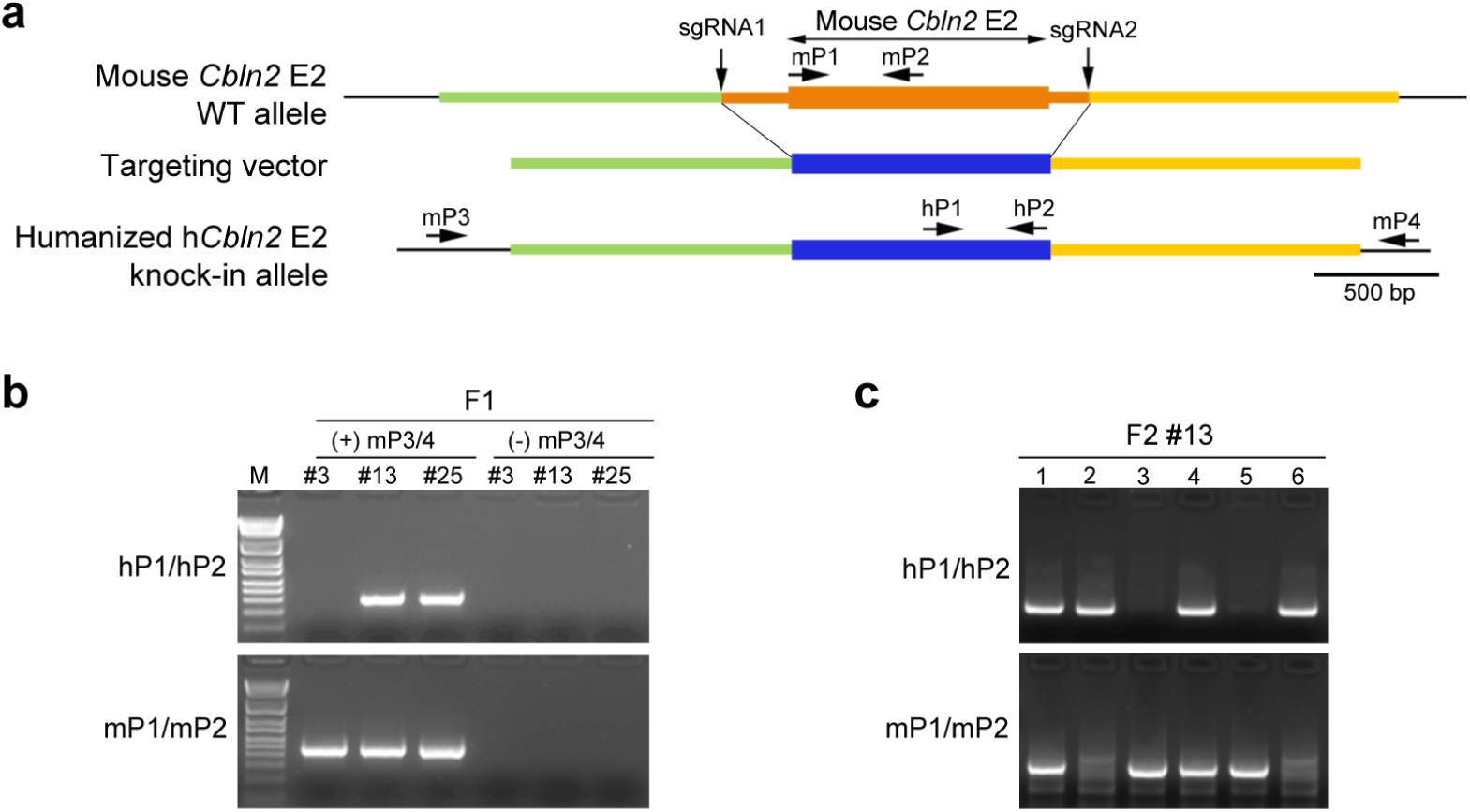
Strategy for generation of humanized *Cbln2* E2 knock-in mice by CRISPR-Cas9 technique. **a**, Positions of single guide RNAs (sgRNA1 and 2) to introduce double strand breaks in the genomic DNA and targeting vector to replace WT mouse *Cbln2* E2 region with that of human *CBLN2* E2 (humanized h*Cbln2* E2) are shown. **b**, Genotyping strategy for F1 mice. Germ line transmission in the F1 generation was confirmed by nested PCR using the primer set of mP3/mP4, followed by hP1/P2 as indicated in a. Two founders #13, and #25 were obtained. **c**, Mice in the following generation were genotyped by PCR with hP1/P2 and mP1/mP2. An example of genotyping for F2 of line #13 mice is shown.

**Extended Data Figure 8.**
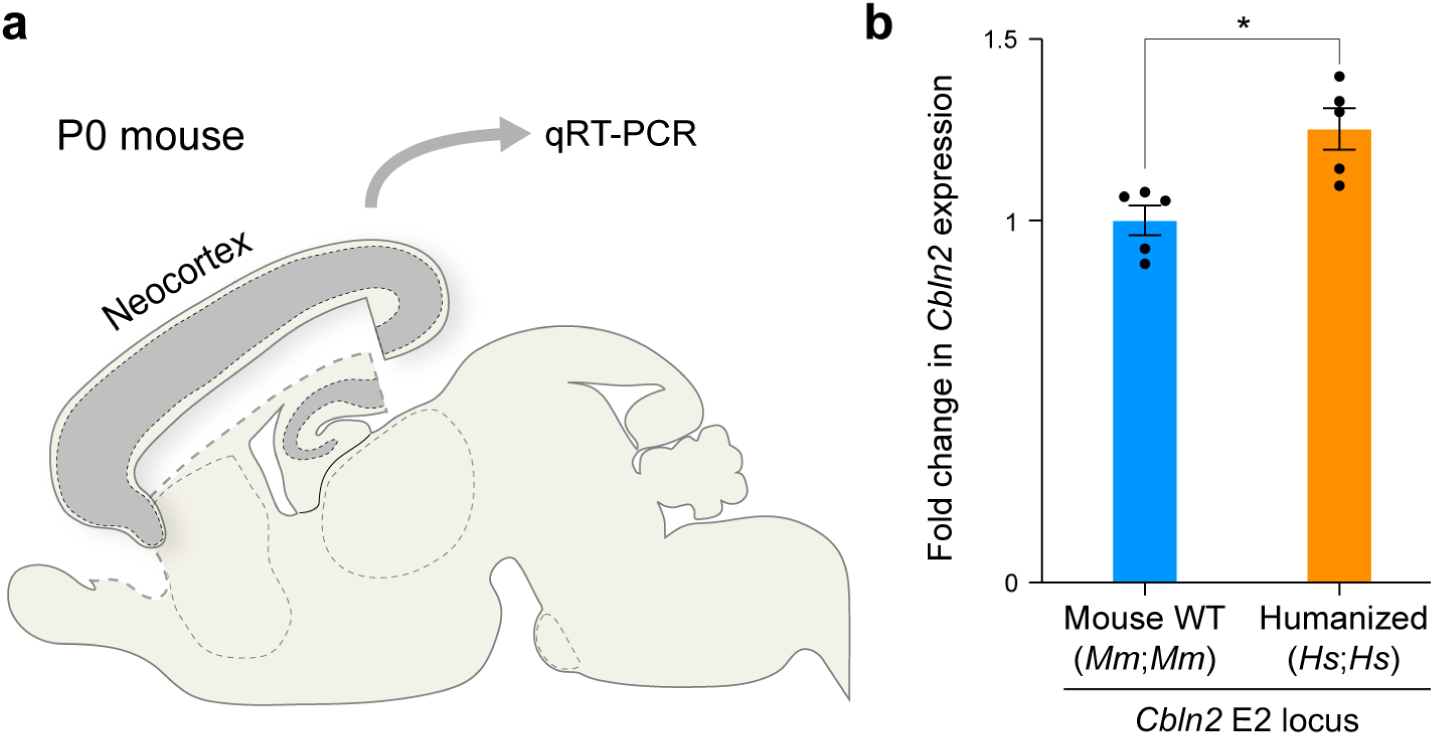
Humanized *Cbln2* E2 knock-in mouse shows increased *Cbln2* expression in neonatal neocortex. **a**, Comparison of *Cbln2* expression between WT *Cbln2* E2 (*Mm;Mm*) and h*Cbln2* E2 (*Hs;Hs*) neocortex at P0 using quantitative reverse transcription-PCR. RNA was extracted from the neocortex following the removal of hippocampus, olfactory bulb, and subpallial regions. Two-tailed Student’s test; *P = 0.007; Error bars: S.E.M.; N = 5 per genotype.

**Extended Data Figure 9.**
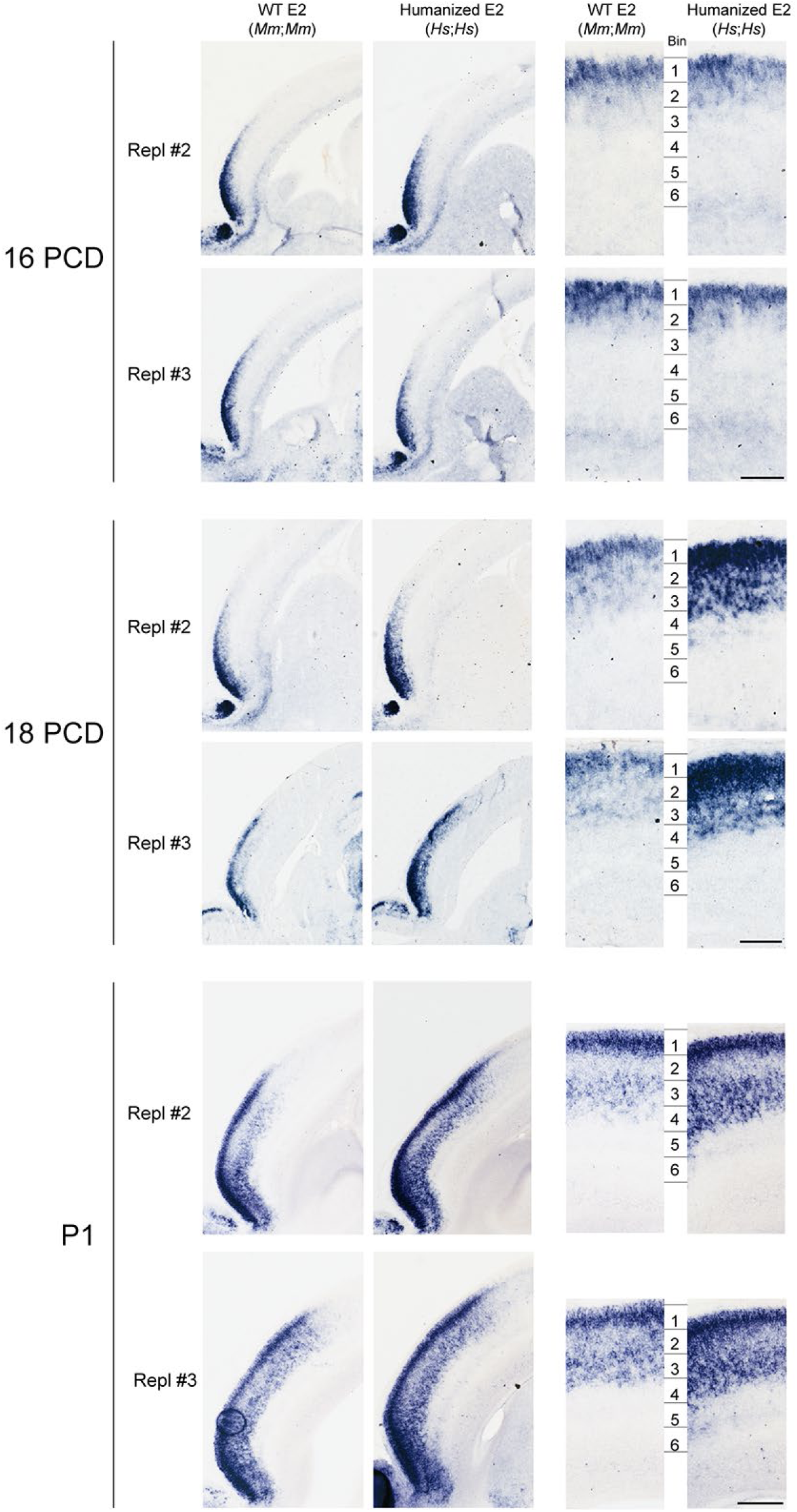
Additional replicates from Figure 3. The neocortex of the humanized h*Cbln2* E2 knock-in prenatal and neonatal mice exhibits upregulated *Cbln2* in both upper and deeper layers, when compared to mice carrying WT *Cbln2* E2. Additional two replicates (Repl #2 and 3) not shown in Figure 3a are shown. Scale bar: 200 μm.

**Extended Data 10.**
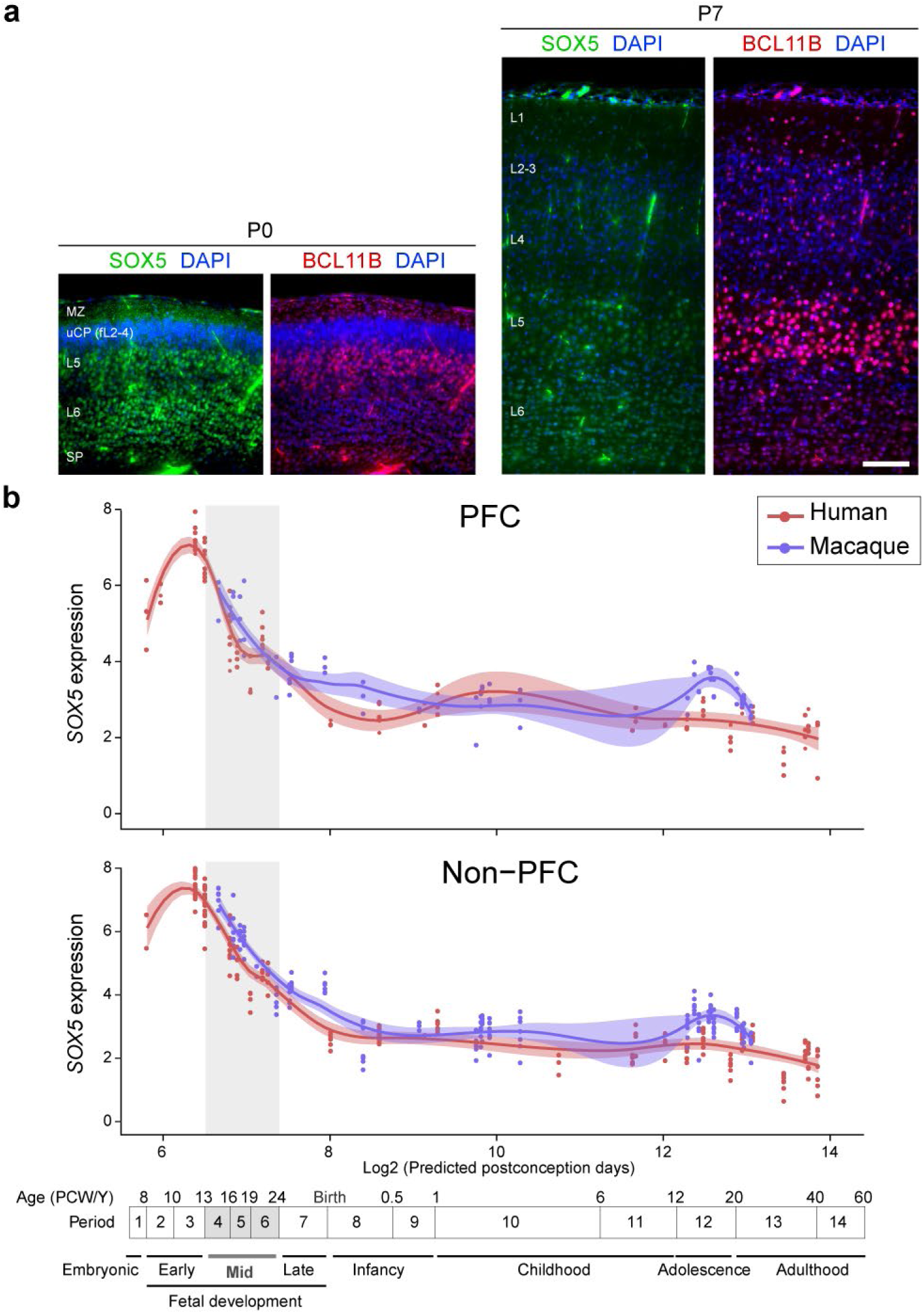
Expression of *Sox5* in the developing mouse and human neocortex. **a**, Double immunofluorescent staining for SOX5 and BCL11B in P0 and P7 mouse neocortex. While the intensity of BCL11B increased at P7 as compared to P0, SOX5 intensity decreased at P7 compared P0. N = 3 per condition. Scale bar: 100 μm. **b**, SOX5 expression in the PFC and non-PFC areas of the cerebral cortex of human and macaque across development. Red and blue lines indicate human and macaque, respectively. Vertical gray box demarcates mid-fetal developmental periods. Timeline of human and macaque development and the associated periods designed by Kang et al. ^40^ shown below.

